# Insulin-Like Growth Factor I Modulates Sleep Through Hypothalamic Orexin Neurons

**DOI:** 10.1101/2020.03.20.000000

**Authors:** Jonathan A. Zegarra-Valdivia, Jaime Pignatelli, Maria Estrella Fernandez de Sevilla, Ana M. Fernandez, Victor Munive, Laura Martinez-Rachadell, Angel Nuñez, Ignacio Torres Aleman

## Abstract

Although metabolic and sleep disturbances are commonly associated, the underlying processes are not yet fully defined. Insulin-like growth factor-I (IGF-I), an anabolic hormone that shows a circadian pattern in the circulation and activity-dependent entrance in the brain, is associated to sleep regulation along evolution. However, its role in this universal homeostatic process remains poorly understood. We now report that the activity of orexin neurons, a discrete cell population in the lateral hypothalamus that is involved in the circadian sleep/wake cycle and arousal, is modulated by circulating IGF-I. Furthermore, mice with blunted IGF-I receptor activity in orexin neurons have lower levels of orexin in the hypothalamus, show altered electrocorticographic patterns with predominant slow wave activity, reduced onset-sleep latency, and less transitions between sleep and awake stages. Collectively, these results extend the role of this pleiotropic growth factor to shaping sleep architecture through regulation of orexin neurons. We speculate that poor sleep quality associated to diverse conditions may be related to disturbed brain IGF-I input to orexin neurons.

## Introduction

Metabolism and sleep are functionally linked ^1,2^. Anabolic signals of the somatotropic axis encompassing pituitary growth hormone (GH), and liver insulin-like growth factors I and II (IGF-I and IGF-II), together with the other components of this axis, the hypothalamic peptides GHRH and somatostatin, have been reported to participate in sleep regulation in various ways. Thus, GH secretion is modulated by sleep ^3^, but without affecting it ^4^. IGF-I, together with GHRH and somatostatin, influence this universal homeostatic process ^5-8^, while little is known of a possible relationship between IGF-II and sleep. Significantly, the link between IGF-I and sleep spans the evolutionary tree ^9-11^, and at least in mammals, sleep modulates circulating IGF-I levels ^12-14^, suggesting a bi-directional relationship between sleep and IGF-I (and probably other members of the somatotropic axis). However, the mechanisms underlying these relationships are still undefined.

In our search for a mechanistic understanding of the relationship of IGF-I with the sleep/wake cycle, we turned our attention to orexin neurons (also termed hypocretin neurons) located in the Perifornical area (PeF) in the lateral hypothalamus ^15,16^. These neurons are key regulators of a wide range of physiological functions, such as the sleep/wake cycle ^17^ or physical activity ^18^, and receive multiple endocrine modulatory inputs, including insulin, a hormone closely related to IGF-I ^19^. Orexin neurons also express IGF binding protein 3, a major regulator of IGF-I bio-availability ^20^, suggesting that they are targeted by this growth factor.

Since orexin neurons are involved in the facilitation and/or maintenance of arousal ^21-23^, we hypothesized that IGF-I may form part of a feedback loop between the periphery and the hypothalamus to control the circadian pattern of activity by modulating orexinergic function. Indeed, circulating IGF-I enters into the brain in response to physical activity ^24^, through an activity-dependent process ^25^. Furthermore, recent evidence indicates that circulating IGF-I shows a circadian pattern and is a circadian entrain cue ^26,27^, affecting hypothalamic clock genes ^28^.

In this work we used in vivo recordings, and genetic inactivation of the IGF-I receptor (IGF-IR) in orexin neurons through the Cre/Lox system, and determined that IGF-I modulates their activity, ultimately shaping sleep architecture. Our results help explain the functional link between IGF-I and the sleep/wake cycle.

## Materials and Methods

### Materials

Antibodies used in this study include rabbit polyclonal c-Fos (Abcam ab190289), rabbit polyclonal IGF-I Receptor-β (Santa Cruz 713/AC) and rabbit anti-IGF-I receptor β XP (Cell Signaling Technology, USA), orexin polyclonal goat antibody (Santa Cruz 8070), orexin polyclonal rabbit antibody (Abcam ab 6214), cre recombinase monoclonal mouse antibody (Millipore clone 2d8), as well as anti-pAkt (Cell Signaling), and monoclonal anti-phosphotyrosine (clone PY20, BD Transduction laboratories, USA). Human recombinant IGF-I was from Pre-Protech (USA).

### Animals

Adult female and male C57BL/6J mice (Harlan Laboratories, Spain), Cre/Lox mice lacking functional IGF-I receptors in orexin neurons (Floxed IGF-IR/Orexin Cre: Firoc mice), and channelrhodopsin (ChR)/Orexin-Cre mice (Ox-ChR mice) were used (3-6 months old; 26 – 33g). Littermates were used as controls. In a subset of electrophysiological experiments, wild type mice were included in the control group since no differences were seen with littermates (see results). The estrous cycle of the female mice was not controlled. Experiments were done during the light phase, except when indicated.

Firoc mice were obtained by crossing Orexin-Cre mice (a kind gift of T Sakurai, Tsukuba Univ, Japan ^29^ with IGF-IR^f/f^ mice (B6, 129 background; Jackson Labs; stock number: 012251; exon 3 floxed). Cre recombination of the floxed IGF-IR results in deletion of exon 3, functional inactivation of the receptor ^30^, and substantial reduction of IGF-IR expression ^31^. ChR mice were obtained from L Menendez de la Prida (Cajal Institute) and crossed with Orexin-Cre mice. Genotyping of Firoc mice was performed using the primers: 5’-GGT TCG TTC ACT CAT GGA AAA TAG-3’ and 5’-GGTATCTCTGACCAG AGTCATCCT-3’ for Orexin-Cre and: 5’-CTT CCCAGCTTGCTACTCTAGG-3’and 5’-AGGCTTGCAATGAGACATGGG-3’ for IGF-IR^f/f^. Ox-Chr mice were genotyped using the same Orexin-Cre primers, ChR primers: 5’-GCATTAAAGCAGCGTATCC-3’and 5’-CTGTTCCTGTACGGCATGG-3’, and wild type primers: 5’-AAGGGAGCT GCAGTGGAGTA-3’, and 5’-CCGAAAATCTGTGGGAAGTC-3’. We confirmed Cre-mediated recombination in the IGF-IR locus between the two sequences flanking the IGF-IR exon-3 only in the hypothalamus of Firoc mice using the previously described primer pair P3 (5‘-TGAGACGTAGGGAGATT GCTGTA-3’) and P2 (5’-CAGGCTTGCAA TGAGACATGGG-3’) ^32^. These primers amplify a fragment of 320bp only when recombination has occurred. Further, in situ hybridization using RNAscope (2.5 HD Detection kit – Red; #322350; ACD, USA) with an IGF-IR exon 3-specific probe combined with immunocytochemistry with anti-orexin antibodies was also performed to confirm deletion of exon 3 in orexin cells.

DNA from brain tissue was isolated using Trizol Reagent and ethanol precipitation. 10 ng of genomic DNA was used in a PCR reaction containing 1X reaction buffer, 1 nM of each primer, 0.2 mM of dNTPS and 0.75 µl of DFS-Taq DNA polymerase (Bioron, GmbH). The thermocycler program was 92°C, 3 min and 30 cycles of 94°C, 30 sec; 65°C, 30 sec; 72°C, 30 sec, and after that, a final extension step at 72°C for 2 min was performed. Amplicons were analyzed in 3% agarose gels stained with SYBRsafe (Thermofisher).

Animals were housed in standard cages (48 × 26 cm^2^, 5 per cage), and kept in a room with controlled temperature (22°C) under a 12-12h light-dark cycle. Mice were fed with a pellet rodent diet and water *ad libitum*. Animal procedures followed European guidelines (2010/63, European Council Directives) and were approved by the local Bioethics Committee (Government of the Community of Madrid).

### Administration of IGF-I

IGF-I was dissolved in saline and intraperitoneally (ip) injected (1µg/g body weight). In some experiments, mice were processed for immunocytochemistry (see below) one or two hours after ip IGF-I injection to allow expression of phospho-Akt or c-fos, respectively, whereas in other experiments PeF recordings were carried out immediately after ip injection. Alternatively, IGF-I was locally delivered in the PeF (10 nM; 0.1 μl; coordinates from Bregma: A, −1.95; L, 1 and depth, 4.0 - 4.5mm), and injected animals were thereafter submitted to electrophysiological recordings as described below. Doses were selected based on previous work using pharmacological injections in both systemic and local/ intraparenchymal administration.

### Experimental Design

#### Recordings in anesthetized animals

Mice were anesthetized with isofluorane (2% induction; 1–1.5% in oxygen, maintenance doses), placed in a David Kopf stereotaxic apparatus (Tujunga, CA, USA) in which surgical procedures and recordings were performed, with a warming pad (Gaymar T/Pump, USA) set at 37°C. Local anesthetic (lidocaine 1%) was applied to all skin incisions and pressure points. An incision was made exposing the skull, and small holes were drilled in the skull. Tungsten macroelectrodes (<1 MΩ World Precision Instruments, USA) were used to record the local field potential and the evoked potential in the PeF (coordinates from Bregma: A, −1.95; L, 1 and depth, 4.0 - 4.5mm). Recordings were filtered (0.3–50 Hz), and amplified via an AC preamplifier (DAM80; World Precision Instruments). The LC was stimulated using 120 µm diameter stainless steel bipolar electrodes (World Precision Instruments, coordinates from Bregma: A, −5.4; L, 1 and depth, 4.0 - 4.5mm). Electrical stimulation was carried out with single square pulses (0.3 ms duration and 20-50 µA intensity, delivered at 1 Hz; Cibertec Stimulator, Spain). After a basal recording, insulin or IGF-I were injected locally (Firoc= 13, control= 20) or systemically (Firoc= 10, control= 10).

#### Optogenetics

Optogenetic experiments were performed to identify orexin neurons through light activation using Ox-ChR mice (see Figure 5). Animals were anesthetized with isofluorane, positioned in the stereotaxic apparatus and handled as above. The sagittal midline of the scalp was sectioned and retracted, and a small craniotomy was drilled over the perifornical (PeF) hypothalamic area (same stereotaxic coordinates as above). Optical stimulation of ChR-expressing neurons was achieved with light-emitting diodes (LED; Thomas Recording, Germany) delivered through an optrode, composed of a tungsten microelectrode of 0.5–0.8 MΩ attached to an optical fiber (core diameter 120 µm; Thomas Recording). Unit recordings were performed through the optodre, filtered (0.3–3 KHz), and amplified using a DAM80 preamplifier (World Precision Instruments). Single-unit activity was extracted with the aid of Spike2 software (Cambridge Electronic Design, UK) and sampled at 10 KHz via an analog-to-digital converter built into the Power 1401 data acquisition unit, and fed into a PC computer for off-line analysis with Spike 2 software. The LED was triggered with a square-step voltage command. Stimulation was applied by a single long-lasting pulse of 473 nm light (16 stimuli with a duration of 300 ms) with an illumination intensity of <30mW/mm^2^, which is below the damage threshold of ∼100 mW/mm^2^ for blue light ^33^. The stimulation area was very restricted since total transmitted light power was reduced by 50%, after passing through 100 µm of neuronal tissue, and by 90% at 1 mm ^34^. When LED stimulation was applied to the cortex of Ox-ChR mice, no stimulation was observed.

**Figure 1:**
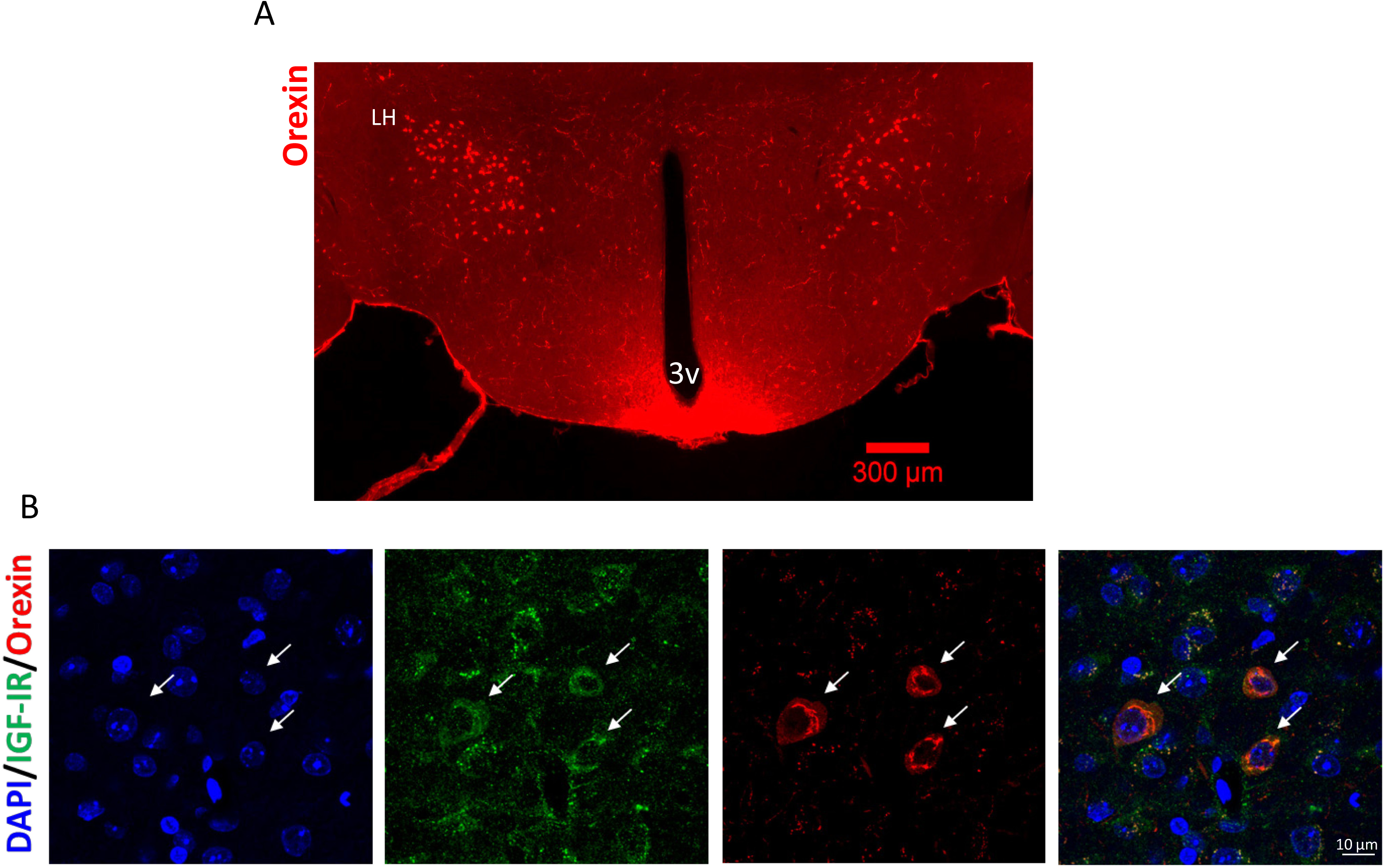
Orexin neurons express IGF-I receptors. **A**, Representative photomicrograph of the mouse hypothalamus showing staining of orexin neurons (red) in the PeF area of the lateral hypothalamus (LH). Note the presence of abundant orexin fibers in the median eminence, as reported by others ^66^. 3v: third ventricle. **B**, Double immunocytochemistry in lateral hypothalamic sections at higher magnification show that orexin neurons (red) express IGF-IR (green). Note that many IGF-IR^+^ cells are not orexinergic, whereas not all hypothalamic cells (DAPI^+^, blue) express IGF-IR. Arrows indicate double stained orexin/IGF-IR cells.

**Figure 2:**
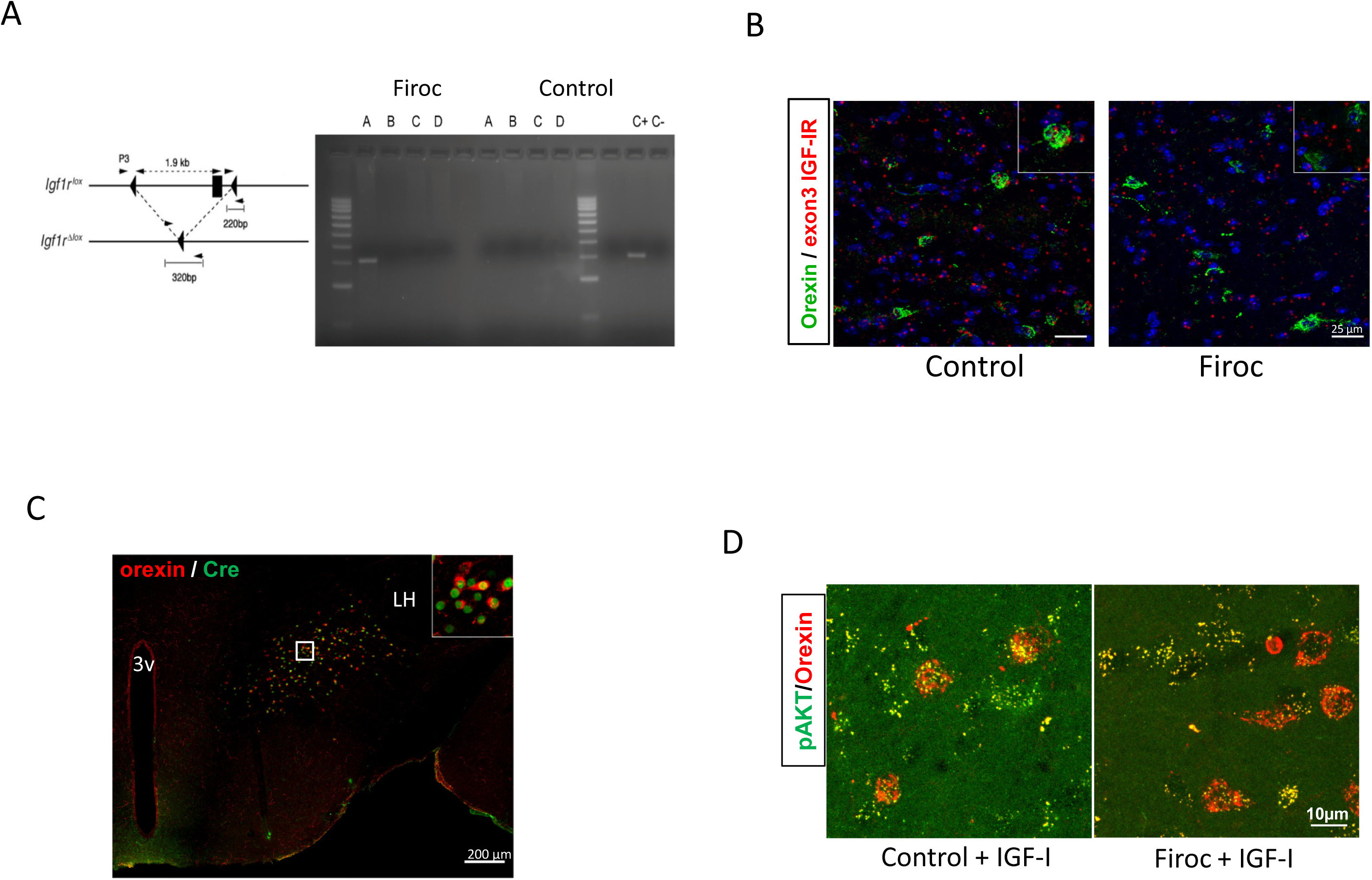
Characterization of mice expressing and inactive IGF-IR in orexin neurons (Firoc mice). **A**, Cre recombination in the IGF-IR gene was analyzed in different brain areas of Firoc and control littermates by PCR using primers P3 and P2 that amplify a 320bp fragment (left diagram) in those cells where Cre recombination has taken place in exon 3 of the IGF-IR. A: hypothalamus, B: ventral thalamus, C: dorsal thalamus, D: hippocampus. Signal is seen only in the hypothalamus of Firoc mice. C^+^: positive control; C^-^: negative control. **B**, Only control littermates, but not Firoc mice, express the non-truncated IGF-IR (wild type) in orexin cells, as determined by RNAScope using an exon 3 IGF-IR-specific probe. Right upper corner: magnification showing the absence of IGF-IR exon-3 mRNA (red) in orexin cells (green). **C**, Double immunocytochemistry confirmed that Firoc mice express Cre (green) in orexin neurons (red). **D**, Systemic injection of IGF-I (1µg/g, ip) stimulates Akt in orexin neurons (red), as determined by immunoreactivity of phosphorylated Akt (pAkt, green) of control littermates, while in Firoc mice co-staining of pAkt and orexin (yellow), is very low. Representative micrographs are shown.

**Figure 3:**
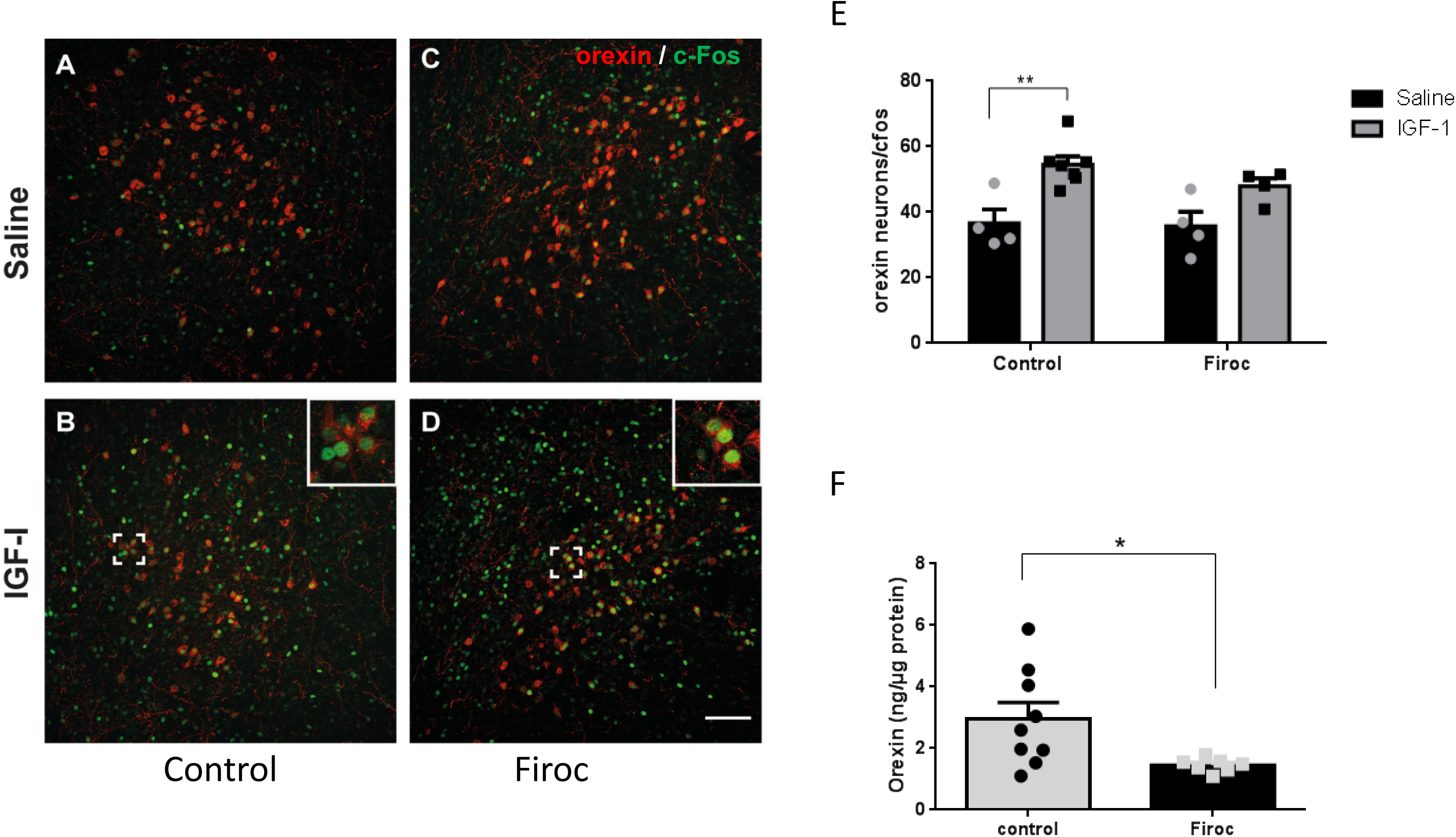
IGF-I stimulates orexin neurons. **A-D**, Double immunocytochemistry of orexin and c-fos in mice injected ip with IGF-I or saline (1µg/g). Double-stained orexin/c-fos cells are shown in the upper right white square in panel B and D, at higher magnification. **E**, Quantification of orexin^+^/c-fos^+^ cells show that their number was significantly increased by IGF-I only in control littermates (*p<0.01; control= 4-7/group, Mann Whitney U Test), but not in Firoc mice (*p<0.05; Firoc= 4-4/group, Mann Whitney U Test). **F**, Levels of orexin in the hypothalamus are significantly decreased in Firoc mice (*p<0.05; Firoc=7, control=9; sex balanced; Unpaired t test).

**Figure 4:**
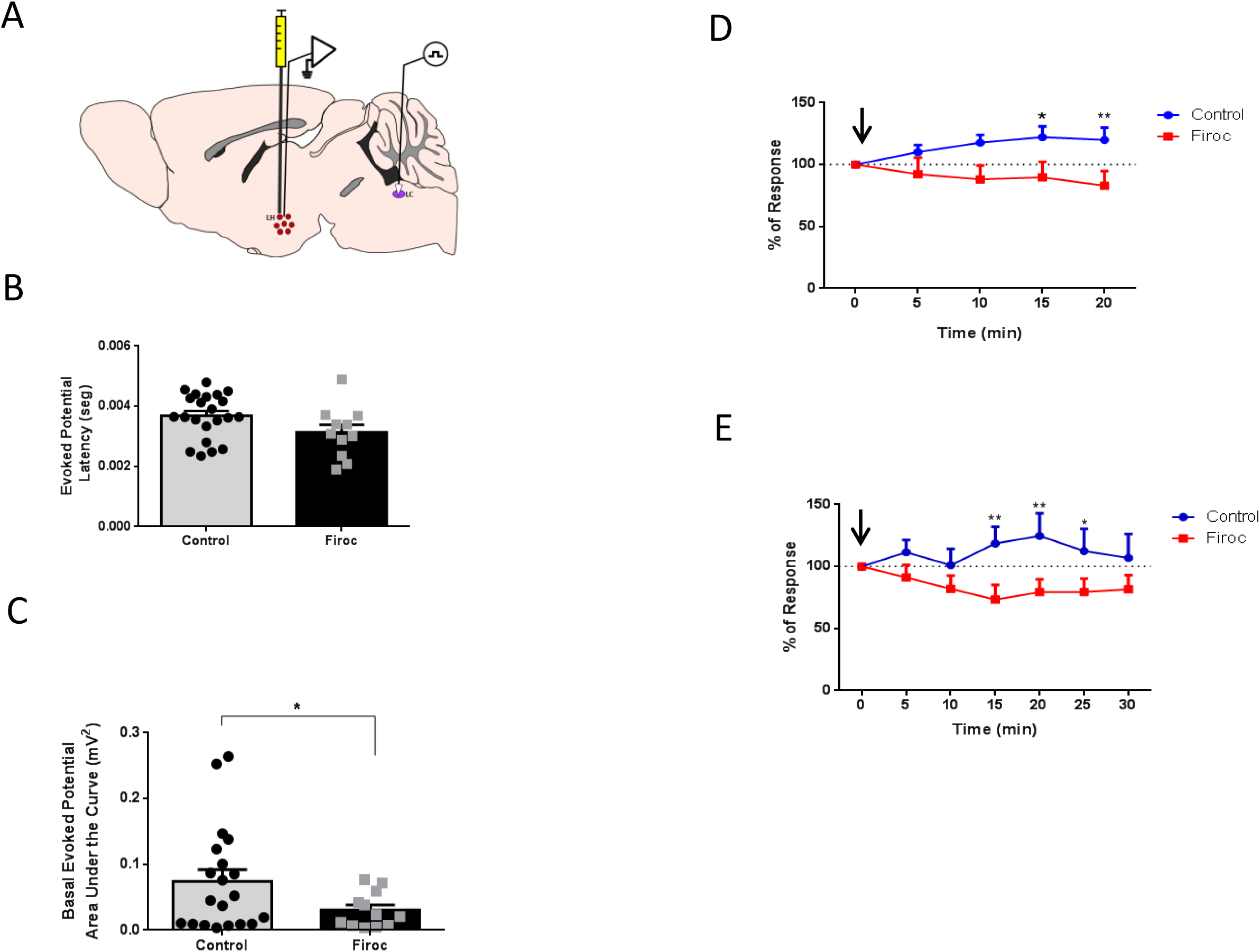
Responses of PeF neurons to Locus Coeruleus (LC) stimulation. **A**, Diagram of experimental design is shown. A stimulating electrode was placed in the LC, and a recording electrode in the PeF at the lateral hypothalamus (LH). **B**, No differences were seen in the latency of the evoked potential in control (a mixed group of wild type mice and littermates), and Firoc mice after stimulation of the LC (Firoc=10, control=19; sex balanced). **C**, Area under the curve of the basal evoked potential after LC stimulation was significantly greater in control than in Firoc mice (*p=0.0315; control=22, Firoc=11; sex balanced, Unpaired t test and Welch’s correction). **D**, Control, but not Firoc mice, responded to local application of IGF-I in the PeF (arrow, 10 μM, 0.1 μl) after LC stimulation. Time course showing the evoked potential of orthodromic impulses after electrical stimulation of the LC in both experimental groups expressed as percentage of basal responses at time 0 (***p< 0.001; control=22, Firoc=11; sex balanced, Two-Way ANOVA, Sidak’s Multiple comparison test). **E**, Intraperitoneal injection of IGF-I (arrow, 1μg/g) increased neuronal activity in PeF orexin neurons after LC stimulation in control (n=10; at 15min **p=0.0011, 20min **p=0.0011, and 25min *p=0.0301), but not in Firoc mice (n=10, sex balanced, Two-Way ANOVA, Sidak’s Multiple comparison test).

**Figure 5:**
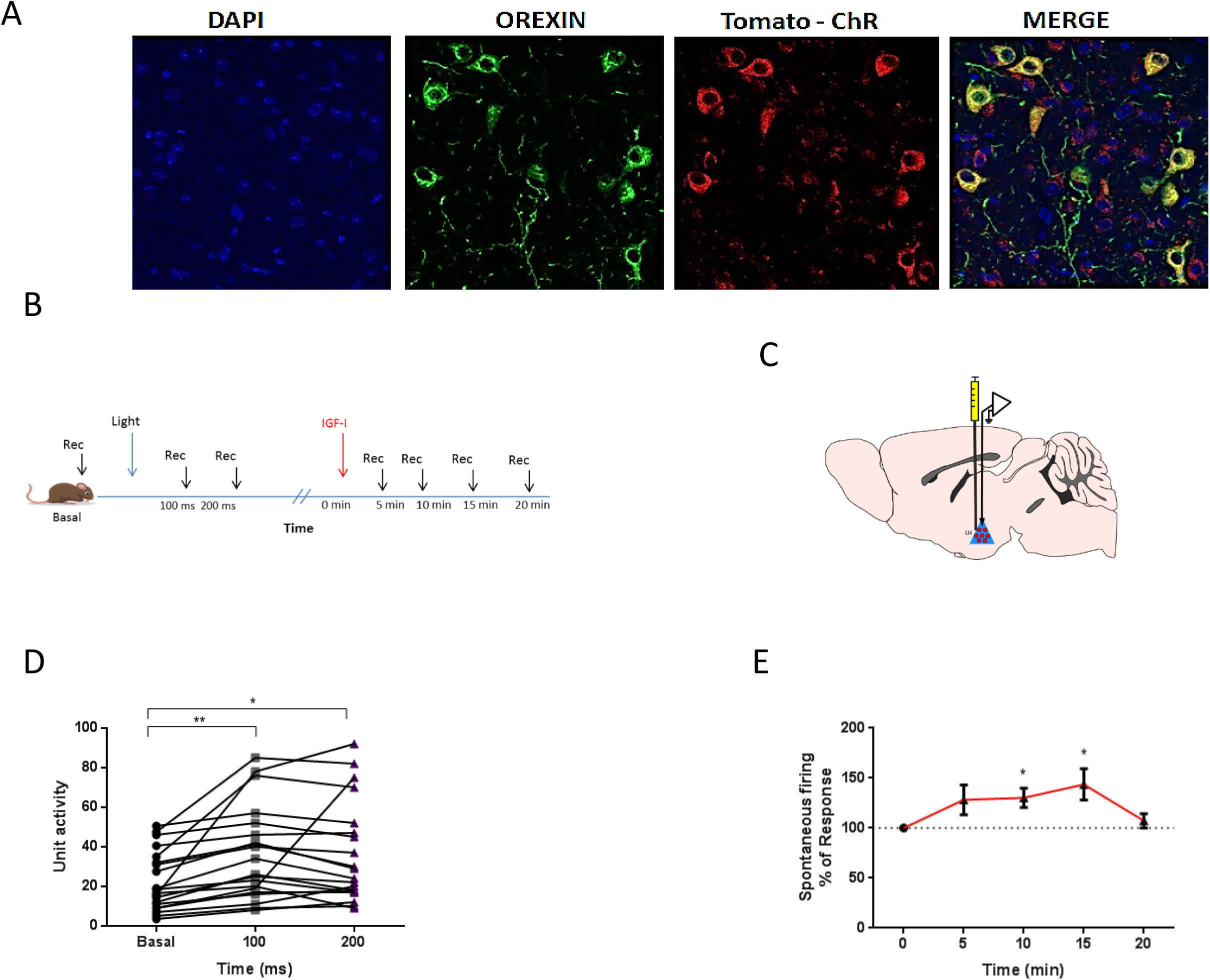
Orexin neurons are activated by IGF-I. **A**, Representative micrographs of orexin neurons (green) co-labeled with Tomato-ChR (red) in Ox-ChR mice. DAPI staining of cell nuclei is also shown. **B**, Diagram of experimental design used in optogenetic identification of orexin neurons. After recording (Rec) basal electrophysiological activity in the PeF area, light stimulation was delivered and recordings continued thereafter. Once active neurons were identified, IGF-I was administered and recordings continued for up to 20 minutes more. **C**, Diagram of the positioning of the optrode and the cannula in the LH in optogenetic experiments. **D**, Unit recordings in the PeF area under optogenetic activation for a total of 16 light stimuli. Basal activity is the mean of unit activity between −200ms and −100ms. Values at 100ms, and 200ms represent unit activity during blue light stimulation (**p=0.0037; n=20-20-20/group; ***p<0.001; Basal vs 100ms:**p=0.002; Basal vs 200ms: *p=0.0320; 100ms vs 200ms: p=0.999; sex balanced, Repeated Measure One-Way ANOVA, Tukey’s Multiple comparison test). **E**, Local injection of IGF-I also stimulated the spontaneous activity of optogenetically identified orexin neurons in the PeF area at 10min (*p=0.0261), and 15min (*p=0.0121). Ox-ChR mice=16; sex balanced, Kruskal-Wallis test and Dunn’s Correction test.

#### Electrocorticogram (ECG) recordings in freely moving animals

Mice were anesthetized as indicated above, and placed in a stereotaxic device. The skin was cut along midline and a craniotomy made (0.5 mm diameter) in the area of the primary somatosensory area (S1). A stainless-steel macro-electrode of <0.5 MΩ was placed without disrupting the meninges to register the cortical electrical activity (ECG), using a DSI Implantable Telemetry device (Data Sciences International, USA). After surgery, mice remain in their cages a minimum of 4 days to recover. ECG was registered for 60 minutes in two days (30 min per day, from 15, 00 – 18, 00). Signals were stored in a PC using DSI software and filtered off-line between 0.3–50 Hz with Spike 2 software. ECG segments of 5 minutes were analyzed with this software using the Fast Fourier Transform algorithm to obtain the power spectra. The mean power density was calculated for 5 different frequency bands that constitute the global ECG: delta band (0.3–5 Hz), theta band (5–8 Hz), alpha band (8–12 Hz), beta band (12–30 Hz), and gamma band (30–50 Hz). The total power of the five frequency bands was considered 100%, and the percentage of each frequency band was calculated for the 60 minutes. To determine the global ECG during the active phase we used the following animals: Firoc= 5, control= 5; during the passive phase: Firoc= 7, control= 12, and for the sleep pattern in the passive phase: Firoc= 7, control= 6.

To assess sleep/awake status, segments of 30 sec of the ECG recording were analyzed according to the presence of slow waves (0.3–5 Hz), fast waves (>12 Hz), and mouse’s movements. Every segment with an equal quantity of slow/fast waves was considered a transition phase. The total % of events (sleep, transitions, or awakening state) was measured (Firoc= 6, control= 9). We measured the latency to sleep onset taking these periods (Firoc= 6, control= 9). ECG recording were performed at different time points through a remote computer.

### Data Analysis

Evoked potentials elicited by LC electrical stimulation (20-50 μA; 0.3 ms duration; at 1 Hz) were calculated. The peak latency was calculated as time elapsed between the stimulus onset and the peak of the second evoked potential wave (orthodromic response, with a latency of 3.5 ± 0.81 ms). To quantify the orthodromic response, the area under the curve of the positive wave was measured from the beginning of the positive slope. On the other hand, single-unit responses were measured from the peri-stimulus time histogram (PSTH; 1ms bin width; 16 stimuli) as the number of spikes evoked during the 0–100 ms or 100-200 ms time-windows after stimulus onset (blue-light pulse). Plots of the unit activity show the percentage of variation respect to basal period (5 mins). In optogenetic experiments, outliers and recordings that did not elicit at least a 70% of increment activity were removed from the analysis to minimize interference from non-orexinergic neurons.

### Immunoassays

#### Immunocytochemistry

Animals were deeply anesthetized with pentobarbital (50 mg/kg) and perfused transcardially with saline 0.9% and then 4% paraformaldehyde in 0.1 M phosphate buffer, pH 7.4 (PB). Coronal 50-μm-thick brain sections were cut in a vibratome and collected in PB 0.1 N. Sections were incubated in permeabilization solution (PB 0.1N, Triton X-100, NHS 10%), followed by 48 hours incubation at 4°C with primary antibody (1:500) in blocking solution (PB 0.1N, Triton X-100, NHS 10%). After washing three times in 0.1 PB, Alexa-coupled goat/rabbit polyclonal secondary antibodies (1:1000, Molecular Probes, USA) were used. Finally, a 1:1000 dilution in PB of Hoechst 33342 was added for 3 minutes. Slices were rinsed several times in PB, mounted with gerbatol mounting medium, and allowed to dry. Omission of primary antibody was used as control. Confocal analysis was performed in a Leica (Germany) microscope. For double stained orexin/c-fos counting, 4 sections per animal were scored using the Imaris software, as described ^35^.

#### ELISA

Hypothalamus from control and Firoc mice were dissected on ice and immediately frozen on dry ice and kept at −80°C until use. Orexin concentration in hypothalamus was determined using Orexin-A ELISA (Phoenix peptides, Inc). Peptide extraction was performed following the manufacturer’s instructions. Briefly, tissue was homogenized in 1 N acetic buffer and boiled 20 minutes at 100°C. Samples were then filtered through a Sep-pak® C-18 cartridge (Sep-Pak®, Millipore, USA), eluted in 2ml of methanol and dried using centrifugal concentration (SpeedVac). Each sample was rehydrated in 150µl of ELISA buffer. Samples were assayed in duplicate. Values were normalized with respect to the total protein on each sample determined by the BCA method (Sigma-Aldrich).

### Statistical Analysis

Statistical analysis was performed using GraphPad Prism 6 software (San Diego, CA, USA), and R Package (Vienna, Austria). Depending on the number of independent variables, normally distributed data (Kolmogorov-Smirnov normality test), and the experimental groups compared, we used either Student’s t-test or two-way ANOVAs followed by Sidak’s multiple comparison test. For non-normally distributed data, we used the Mann Whitney U test for comparing two groups, Kruskall-Wallis or Friedman test, with Dunn’s multiple comparisons as a *Post Hoc* analysis, as well as Scheirer-Ray Test, a non-parametric alternative to multi-factorial ANOVA. Sample size for each experiment was chosen based on previous experience and aimed to detect at least a p<0.05 in the different tests applied, considering a reduced use of animals. Results are shown as mean ± standard error (SEM) and *p* values coded as follows: *p< 0.05, **p< 0.01, ***p< 0.001. Animals were included in each experimental group randomly by the researcher, without blinding.

## Results

### Inactivation of IGF-IR in orexin neurons

Since orexin neurons express IGF-I receptors (Figure 1), we inactivated them to determine their role in orexinergic function. We genetically ablated IGF-IR activity using the Cre/Lox system following previous published procedures ^36^. Mice with inactive IGF-IR in orexin neurons (Firoc mice) express a truncated IGF-IR in the hypothalamus, as determined by PCR detection of the truncated IGF-IR (Figure 2A). Within the hypothalamus, only orexin neurons did not express the native IGF-IR, as determined by in situ hybridization using an RNA probe that detects the intact IGF-IR combined with orexin immucytochemistry (Figure 2B). To confirm Cre-mediated recombination in orexin neurons, double orexin/Cre immunocytochemical staining in Firoc mice show that the majority of cells expressing Cre were orexin neurons (Figure 2C). Further, orexin responses to IGF-I were blunted in Firoc mice, as systemic injection of IGF-I resulted in Akt phosphorylation, a kinase downstream of IGF-IR, in littermate control mice but not in Firoc (Figure 2D).

To confirm functional inactivation of IGF-IR in orexin neurons of Firoc mice, we injected IGF-I (1μg/g, ip), and two hours later we determined the number of orexin neurons expressing c-fos. In control mice, IGF-I significantly increased the number of orexin neurons expressing c-fos (double labeled orexin^+^/c-fos^+^ cells) as compared to saline injected mice (Control: 4-7/group, saline: 36.61 ± 8.397; IGF-I: 54.49 ± 6.638, **p= 0.0076; see Figure 3A, B, E). In Firoc mice a smaller, non-significant increase was seen (Firoc: 4-4/group, saline: 35.68 ± 8.825; IGF-I: 47.89 ± 4.859, p= 0.1679; see Figure 3C,D,E), indicating reduced sensitivity to IGF-I in orexin neurons. The increment in c-fos labeling of orexin neurons seen in Firoc mice after IGF-I administration probably is due to the stimulation by IGF-I of excitatory afferents to orexin neurons and/or the fact that Cre/Lox recombination of IGF-IR in mutant mice did not occur in 100% of orexin neurons (Figure 2B). Furthermore, Firoc mice had significantly lower levels of orexin in the hypothalamus (Firoc= 7, 1.45 ± 0.22; Control= 9, 2.96 ± 1.57, *p= 0.0211; see Figure 3F).

### IGF-I modulates the activity of orexin neurons

Since IGF-I significantly increases the number of orexin neurons expressing c-fos, a marker of cell activation, we determined regulation of the activity of orexinergic neurons by IGF-I using electrophysiological recordings. We studied the activation of PeF neurons to locus coeruleus (LC) stimulation in Firoc mice and control littermates (Figure 4A). The LC, a major connection of orexin neurons ^37^, provides feedback information to PeF neurons by facilitating its firing activity ^38^. Electrical stimulation delivered in the LC (20-50 μA; 0.3 ms duration; at 1 Hz) evoked a fast orthodromic potential in the PeF region. The orthodromic potential consisted of a positive wave with similar mean peak latency in both groups: 3.7± 0.7ms in control littermates (n= 22), and 3.1± 0.8 ms in Firoc mice (n= 11; p=0.08). This indicates that the LC input to the PeF is unaffected in Firoc mice.

However, while the latency of the evoked potential was normal in Firoc mice (Figure 4B), when we calculated its area to quantify changes in the amplitude and duration of PeF responses to LC stimulation, a decrease was observed in Firoc mice. LC stimulation evoked a response area of 74.6±11.9 μV^2^ (n= 22) in control mice, and a significantly lower in Firoc mice: 30.6±2.7 μV^2^ (n= 11; *p= 0.0315; Figure 4C). Importantly, local injection of IGF-I (10 nM; 0.1 μl) in the PeF region of control mice increased the response to LC stimulation (n= 22; ***p< 0.001; Figure 4D). In contrast, Firoc mice (n= 11) were not affected by local injection of IGF-I in the PeF area (Figure 4D). To determine whether systemic IGF-I can also stimulate PeF responses to LC stimulation, we injected ip IGF-I (1µg/g). While in Firoc mice ip IGF-I did not affect LC responses in the PeF area (n=10), in control animals ip IGF-I significantly increased the response area (n=10; *p<0.05, Figure 4E). IGF-I-evoked facilitation in control mice was specific for this growth factor because local application of a same dose of insulin (10 μM; 0.1μl), a closely related hormone, did not facilitate an LC-evoked response (Supplementary Figure).

A second approach to specifically determine whether orexin neurons are activated by IGF-I consisted in optogenetically identify orexin neurons using a mouse expressing channelrhodopsin (ChR) specifically in these cells (Ox-ChR mice, Figure 5A). A blue-light pulse (300 ms duration) able to stimulate a small volume of tissue (about 100–200 µm in radius ^34^) was delivered to the PeF area of Ox-ChR mice using an optrode to perform unit recordings simultaneously with optical stimulation in the same place (Figure 5B,C). Neuronal activity was measured at 100 ms time intervals after light onset (0-100 ms and 100-200 ms), and was compared with the previous 100 ms time interval before time onset (basal condition). Short-lasting blue LED stimuli induced spike firing in neurons of Ox-ChR mice at 0-100ms (36.25 ± 5.25 spikes/100ms) vs basal conditions (22.53 ± 3.37 spikes/100ms; n=20-20/group; **p<0.0022), as well as at 100-200ms (36.30 ± 5.71 spikes/100ms; 20-20/group; *p<0.0320, Figure 5D). The effect lasted 200 ms, recovering baseline activity later, even though the blue-light pulse lasted 300ms. After orexin neurons were identified by their responses to the blue light, local application of IGF-I (10 nM; 0.1 μl) induced a 30.8% increase of their spontaneous firing rate at 10 min (SEM ± 9.56% spikes/100 ms; n=12; *p=0.026), and of 43.6% at 15 min (SEM ± 15.54% spikes/100ms; n=14; *p<0.012), recovering basal levels 20 min after (Figure 5E).

### IGF-I modulates sleep through orexin neurons

The above results, coupled with the evidence that orexin levels may be regulated by IGF-I (Figure 3F), suggested that this growth factor stimulates the activity of orexinergic neurons. Since IGF-I shows a circadian cycle ^39^, and activity-dependent entrance into the brain ^25^, and orexin neurons contribute to the facilitation and/or maintenance of arousal ^40^ in a close connection with the LC ^37^, we studied if electrocorticogram (ECG) patterns are altered in Firoc mice (Figure 6A). We compared the ECG signature in male Firoc mice and controls by calculating the mean percentage of each frequency band measured during the middle part of the dark period (ZT17-ZT19, n=5-5/group). No differences in ECG patterns were found (Figure 6B). However, during the light phase (ZT8-ZT11), Firoc mice displayed differences in δ and θ frequency bands compared to controls (***p<0.001 and **p<0.01, respectively, Figure 6C), indicating that Firoc mice showed a slower ECG than control animals during the light period (their passive phase). This prompted us to determine in more detail if the lack of IGF-I receptor activity in orexinergic neurons seems to favor a slowdown of the ECG, and probably increased sleep periods. Thus, we measured the sleep-onset latency (time since the animal was placed in the recording place until the appearance of δ waves in the ECG without movements) in Firoc vs control mice. As shown in Figure 6D, the latency was shorter in Firoc mice than in controls (n=11/9 group; **p< 0.0058).

**Figure 6:**
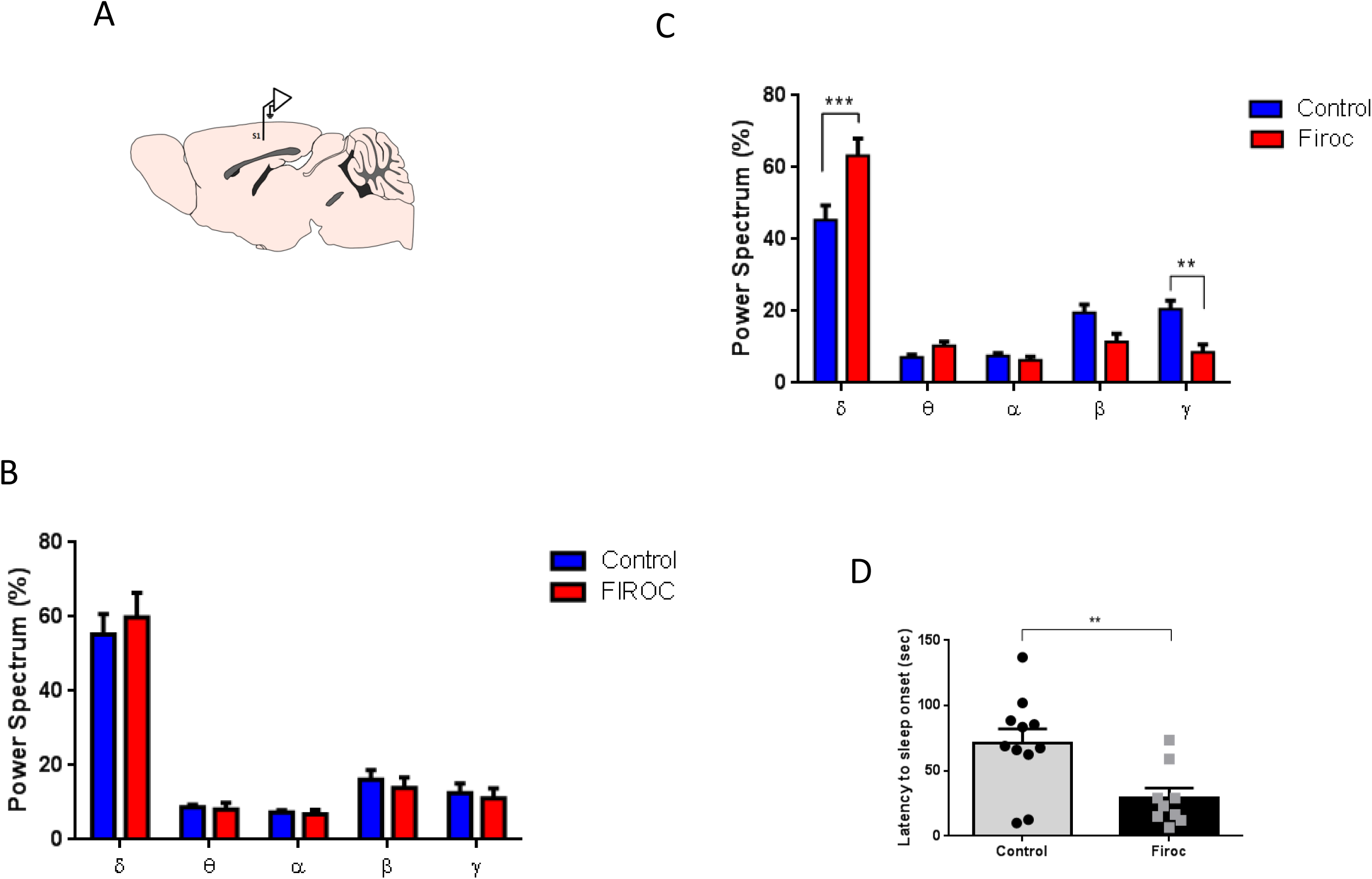
ECG recordings during the light/dark cycle. **A**, Diagram of the intracranial localization of the electrodes in S1cortex; left hemisphere has the reference electrode in all cases. **B**, Power spectra analysis of ECG bands obtained during the dark phase did not show any difference between groups. **C**, ECG analysis during the light phase displays a predominant δ activity together with a reduced γ activity in Firoc mice, as compared to controls (***p<0.001 and **p<0.01; Firoc=7, control=12, males mice only, Two-Way ANOVA, Sidak’s Multiple comparison test).**C**, Latency to sleep-onset was markedly reduced in Firoc mice (**p<0.0058, Firoc=9, control=11, sex balanced, Unpaired t test and Welch’s correction).

Since differences in ECG patterns in Firoc mice were observed along the light period, we analyzed this phase in more detail. Firoc mice showed significantly higher sustained ECG activity in the δ and θ frequency bands during the light phase (ZT3-ZT13; **p<0.01, and ***p<0.001, respectively; n=7-6/group; Figure 7A, B). This slowing of the ECG was accompanied by a significant decrease in α, β, and γ frequency bands (n=7-6/group; *p<0.05, **p<0.01 and ***p<0.001, respectively; Figure 7C-E). Control mice showed alternating awake/sleep periods as evidenced by peaks and valleys in the percentage of δ (Figure 7A), α, β, and γ frequency bands (Figure 7C-E).

**Figure 7:**
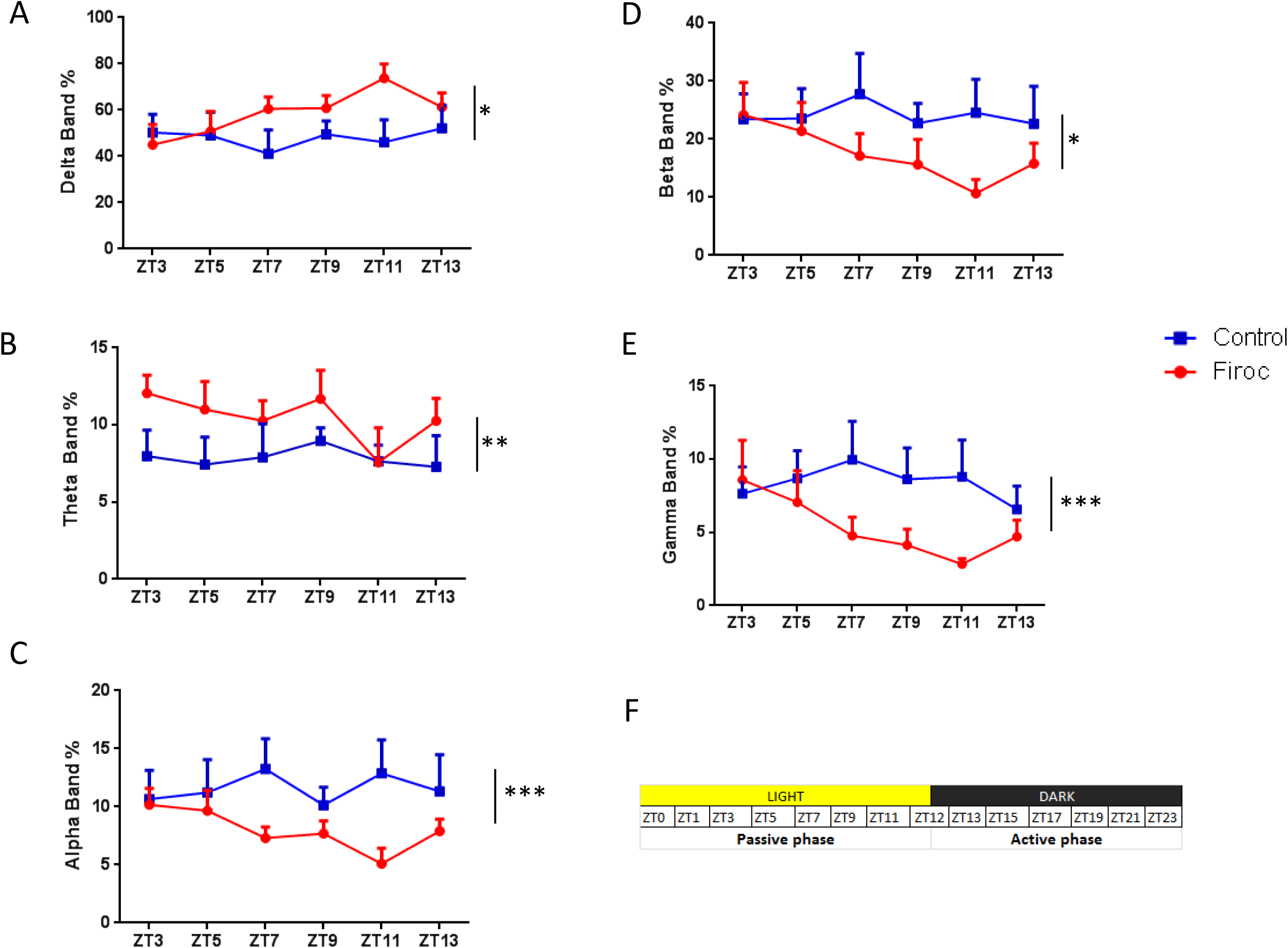
IGF-I shapes sleep architecture through orexin neurons. **A-E**, Sleep architecture during the light phase (ZT 3-13) determined by δ (B), θ (C), α (D), β (E), and γ (F) activity patterns in Firoc (red line), and control mice (blue). An increase in slow activity (δ), and a decrease in fast waves was observed (Firoc=7, control=6; male mice only, Scheirer-Ray Test; *p<0.05, **p<0.01, ***p<0.001). **F**, Chart with the light and dark periods including the corresponding Zeitgeber time.

## Discussion

These results indicate that reduced IGF-I signaling onto orexin neurons results in altered ECG patterns with a predominance of slow waves, suggesting a direct relationship between the anabolic somatotropic axis and sleep through these hypothalamic neurons. Specifically, we show that optogenetically identified PeF neurons are activated by IGF-I -although we cannot entirely rule out that other PeF, non-orexinergic neurons could be activated by IGF-I. However, since blunted IGF-IR activity in the PeF area induces changes in the ECG, as seen in the Firoc mouse, and until now only orexin neurons are known to produce these changes ^38^, we may conclude that most of our results may be explained by a direct activation of orexinergic neurons by IGF-I, facilitating in this way wakefulness. Accordingly, Firoc mice showed an increase in δ waves as well as a decrease in faster ECG activities, especially during the light phase. Thus, IGF-I seems to modulate sleep architecture.

Of note, IGF-I shows a circadian pattern, with highest levels during the sleep period ^27^, when orexin activity should be attenuated ^17^. Thus, it is unlikely that its circadian rhythm will dictate IGF-I signaling to orexin neurons. Rather, it seems more plausible that brain and physical activity ^24,25^, and/or feeding schedule ^26^ will modulate orexin activity, since all these situations are accompanied by increased brain IGF-I input. But more work is needed to determine the specific cues timing IGF-I input to orexin neurons.

While IGF-I is a highly conserved sleep regulatory signal throughout evolution, from invertebrates to vertebrates ^9-11,41^, the mechanisms linking it to sleep-regulatory circuits remain poorly understood. Interestingly, reduced insulin-like signaling in *drosophila* produces also increased sleep ^42^, as we have seen in Firoc mice. Thus, our results provide new insights into this connection in mice, although more studies are needed to clarify whether the functional link between IGF-I and orexin is present in other mammalian species, and even in lower taxa.

Significantly, genetic ablation of orexinergic neurons or mutation of orexin receptors results in a decrease in wakefulness and altered sleep/wake cycles ^43-46^. In turn, lack of IGF-IR activity in orexin neurons results in altered sleep architecture, and reduced sleep-onset latency, consistent with lower orexin levels in these mice. Sleep and wakefulness are mutually exclusive states that cycle with both ultradian and circadian periods. Orexinergic neurons are strongly activated during active wakefulness, decrease their activity during quiet wakefulness, and are silent during the different sleep phases ^47,48^. These neurons are essential for maintaining wakefulness and regulate REM sleep, and their loss results in narcolepsy ^38,43,49,50^. Moreover, orexin neurons are sensitive to metabolic signals ^51^, as well as to peripherally derive circulating factors whose fluctuations may provide information about homeostatic status. Our findings include IGF-I as an additional signal modulating orexin excitability. Thus, it seems that IGF-I input is necessary for orexin neurons to maintain their normal activity. In turn, since altered sleep modifies IGF-I activity, the way sleep influences IGF-I needs to be clarified.

A direct functional connection between IGF-I and orexin may be involved in sleep disturbances such as those that take place during aging ^52^. Old age is related to lower serum IGF-I levels ^53^, impaired brain IGF-I activity ^54^, and reduced number of orexin neurons ^55^. Furthermore, a relationship between sleep disturbances and dementia is widely documented and intensely scrutinized ^56,57^. Sleep disturbances occur very early in the course of Alzheimer’s disease (AD), which is consistent with the finding that brain regions involved in sleep and circadian control are affected early in the pathogenesis of the condition ^58^. On the other hand, long sleep duration in late-life, as well as sleep fragmentation or excessive daytime sleepiness have been associated to increase risk of dementia and poor cognitive performance ^59-63^. Conceivably, impaired IGF-I signaling in AD ^64,65^ may also affect orexinergic function and ultimately, sleep.

This study contains several limitations. First, many of our results are based on a Cre/Lox transgenic mouse that may show ectopic Cre expression, which would result in a contribution of unidentified brain cells expressing IGF-IR in the observed changes. Moreover, Firoc mice show a non-significant increase in c-fos expression in orexin cells after IGF-I that may be related to their stimulation by unidentified IGF-IR-expressing cells and/or orexin cells still expressing fully active IGF-IR. Again, changes observed may also be contributed by these additional cells.

In summary, albeit with the above limitations, our results provide an explanation of the relationship between IGF-I and sleep pointing to orexin neurons as important mediators of this link.

## Acknowledgements

This work was funded by a grant from Ciberned, and from SAF2016-76462 (AEI/FEDER; MINECO). J.A. Zegarra-Valdivia acknowledges the financial support of the National Council of Science, Technology and Technological Innovation (CONCYTEC, Perú) through the National Fund for Scientific and Technological Development (FONDECYT, Perú). We are thankful to M. Garcia for technical support. We also thank the technical assistance of the Cajal Molecular Biology Core Facility (S Fernandez).

## LEGENDS TO FIGURES

**Supplementary Figure:** Local injection of insulin (10 μM; 0.1μl) does not affect neuronal activity in PeF orexin neurons after LC stimulation (Firoc=9, control=17; sex balanced).

## References

1 Arrigoni, E., Chee, M. J. S. & Fuller, P. M. To eat or to sleep: That is a lateral hypothalamic question. Neuropharmacology 154, 34–49, doi:10.1016/j.neuropharm.2018.11.017 (2019).

2 Rutter, J., Reick, M. & McKnight, S. L. Metabolism and the control of circadian rhythms. Annu Rev Biochem 71, 307–331, doi:10.1146/annurev.biochem.71.090501.142857 (2002).

3 Van, C. E., Plat, L. & Copinschi, G. Interrelations between sleep and the somatotropic axis. Sleep 21, 553–566 (1998).

4 Kern, W. et al. Systemic growth hormone does not affect human sleep. J Clin Endocrinol Metab 76, 1428–1432, doi:10.1210/jcem.76.6.8501147 (1993).

5 Steiger, A. et al. Effects of growth hormone-releasing hormone and somatostatin on sleep EEG and nocturnal hormone secretion in male controls. Neuroendocrinology 56, 566–573, doi:10.1159/000126275 (1992).

6 Ziegenbein, M. et al. The somatostatin analogue octreotide impairs sleep and decreases EEG sigma power in young male subjects. Neuropsychopharmacology 29, 146–151, doi:10.1038/sj.npp.1300298 (2004).

7 Obal, F., Jr. et al. Insulin-like growth factor-1 (IGF-1)-induced inhibition of growth hormone secretion is associated with sleep suppression. Brain Res 818, 267–274 (1999).

8 Prinz, P. N. et al. Higher plasma IGF-1 levels are associated with increased delta sleep in healthy older men. J. Gerontol. A Biol Sci. Med Sci 50, M222–M226 (1995).

9 Ashlin, T. G., Blunsom, N. J., Ghosh, M., Cockcroft, S. & Rihel, J. Pitpnc1a Regulates Zebrafish Sleep and Wake Behavior through Modulation of Insulin-like Growth Factor Signaling. Cell Rep 24, 1389–1396, doi:10.1016/j.celrep.2018.07.012 (2018).

10 Skora, S., Mende, F. & Zimmer, M. Energy Scarcity Promotes a Brain-wide Sleep State Modulated by Insulin Signaling in C. elegans. Cell Rep 22, 953–966, doi:10.1016/j.celrep.2017.12.091 (2018).

11 Monyak, R. E. et al. Insulin signaling misregulation underlies circadian and cognitive deficits in a Drosophila fragile X model. Mol Psychiatry 22, 1140–1148, doi:10.1038/mp.2016.51 (2017).

12 Chennaoui, M. et al. Changes of Cerebral and/or Peripheral Adenosine A(1) Receptor and IGF-I Concentrations under Extended Sleep Duration in Rats. Int J Mol Sci 18, doi:10.3390/ijms18112439 (2017).

13 Chennaoui, M., Arnal, P. J., Drogou, C., Sauvet, F. & Gomez-Merino, D. Sleep extension increases IGF-I concentrations before and during sleep deprivation in healthy young men. Appl Physiol Nutr Metab, 1–8, doi:10.1139/apnm-2016-0110 (2016).

14 Monico-Neto, M. et al. REM sleep deprivation impairs muscle regeneration in rats. Growth Factors 35, 12–18, doi:10.1080/08977194.2017.1314277 (2017).

15 de Lecea, L. et al. The hypocretins: hypothalamus-specific peptides with neuroexcitatory activity. Proc Natl Acad Sci U S A 95, 322–327 (1998).

16 Sakurai, T. et al. Orexins and orexin receptors: a family of hypothalamic neuropeptides and G protein-coupled receptors that regulate feeding behavior. Cell 92, 1 page following 696 (1998).

17 Sakurai, T. The neural circuit of orexin (hypocretin): maintaining sleep and wakefulness. Nat Rev Neurosci 8, 171–181 (2007).

18 Kotz, C. M. Integration of feeding and spontaneous physical activity: role for orexin. Physiol Behav 88, 294–301 (2006).

19 Sakurai, T. The role of orexin in motivated behaviours. Nat Rev Neurosci 15, 719–731, doi:10.1038/nrn3837 (2014).

20 Honda, M. et al. IGFBP3 colocalizes with and regulates hypocretin (orexin). PLoS One 4, e4254 (2009).

21 Alam, M. N. et al. Sleep-waking discharge patterns of neurons recorded in the rat perifornical lateral hypothalamic area. J Physiol 538, 619–631, doi:10.1113/jphysiol.2001.012888 (2002).

22 Koyama, Y., Takahashi, K., Kodama, T. & Kayama, Y. State-dependent activity of neurons in the perifornical hypothalamic area during sleep and waking. Neuroscience 119, 1209–1219 (2003).

23 Sakurai, T. et al. Input of orexin/hypocretin neurons revealed by a genetically encoded tracer in mice. Neuron 46, 297–308 (2005).

24 Carro, E., Nunez, A., Busiguina, S. & Torres-Aleman, I. Circulating insulin-like growth factor I mediates effects of exercise on the brain. J. Neurosci 20, 2926–2933 (2000).

25 Nishijima, T. et al. Neuronal activity drives localized blood-brain-barrier transport of serum insulin-like growth factor-I into the CNS. Neuron 67, 834–846 (2010).

26 Crosby, P. et al. Insulin/IGF-1 Drives PERIOD Synthesis to Entrain Circadian Rhythms with Feeding Time. Cell 177, 896–909 e820, doi:10.1016/j.cell.2019.02.017 (2019).

27 Chaudhari, A., Gupta, R., Patel, S., Velingkaar, N. & Kondratov, R. Cryptochromes regulate IGF-1 production and signaling through control of JAK2-dependent STAT5B phosphorylation. Mol Biol Cell 28, 834–842, doi:10.1091/mbc.E16-08-0624 (2017).

28 Breit, A. et al. Insulin-like growth factor-1 acts as a zeitgeber on hypothalamic circadian clock gene expression via glycogen synthase kinase-3beta signalling. J Biol Chem, doi:10.1074/jbc.RA118.004429 (2018).

29 Matsuki, T. et al. Selective loss of GABA(B) receptors in orexin-producing neurons results in disrupted sleep/wakefulness architecture. Proc Natl Acad Sci U S A 106, 4459–4464, doi:10.1073/pnas.0811126106 (2009).

30 Xuan, S. et al. Defective insulin secretion in pancreatic beta cells lacking type 1 IGF receptor. The Journal of clinical investigation 110, 1011–1019, doi:10.1172/JCI15276 (2002).

31 Baker, J., Liu, J. P., Robertson, E. J. & Efstratiadis, A. Role of insulin-like growth factors in embryonic and postnatal growth. Cell 75, 73–82 (1993).

32 Xuan, S. et al. Defective insulin secretion in pancreatic beta cells lacking type 1 IGF receptor. J Clin Invest 110, 1011–1019, doi:10.1172/JCI15276 (2002).

33 Cardin, J. A. et al. Targeted optogenetic stimulation and recording of neurons in vivo using cell-type-specific expression of Channelrhodopsin-2. Nat Protoc 5, 247–254, doi:10.1038/nprot.2009.228 (2010).

34 Aravanis, A. M. et al. An optical neural interface: in vivo control of rodent motor cortex with integrated fiberoptic and optogenetic technology. J Neural Eng 4, S143–156, doi:10.1088/1741-2560/4/3/S02 (2007).

35 Martinez-Rachadell, L. et al. Cell-specific expression of insulin/insulin-like growth factor-I receptor hybrids in the mouse brain. Growth Horm IGF Res 45, 25–30, doi:10.1016/j.ghir.2019.02.003 (2019).

36 Franco, C. et al. A role for astrocytes in cerebellar deficits in frataxin deficiency: Protection by insulin-like growth factor I. Mol Cell Neurosci 80, 100–110, doi:10.1016/j.mcn.2017.02.008 (2017).

37 Carter, M. E. et al. Mechanism for Hypocretin-mediated sleep-to-wake transitions. Proc Natl Acad Sci U S A 109, E2635–2644, doi:10.1073/pnas.1202526109 (2012).

38 Tortorella, S., Rodrigo-Angulo, M. L., Nunez, A. & Garzon, M. Synaptic interactions between perifornical lateral hypothalamic area, locus coeruleus nucleus and the oral pontine reticular nucleus are implicated in the stage succession during sleep-wakefulness cycle. Front Neurosci 7, 216, doi:10.3389/fnins.2013.00216 (2013).

39 Chaudhari, A., Gupta, R., Patel, S., Velingkaar, N. & Kondratov, R. Cryptochromes regulate IGF-1 production and signaling through control JAK2 dependent STAT5B phosphorylation. Mol Biol Cell 28, 834–842, doi:10.1091/mbc.E16-08-0624 (2017).

40 Adamantidis, A. R., Zhang, F., Aravanis, A. M., Deisseroth, K. & de Lecea, L. Neural substrates of awakening probed with optogenetic control of hypocretin neurons. Nature 450, 420–424, doi:10.1038/nature06310 (2007).

41 Zegarra-Valdivia, J. A., Santi, A., Fernandez de Sevilla, M. E., Nunez, A. & Torres Aleman, I. Serum Insulin-Like Growth Factor I Deficiency Associates to Alzheimer’s Disease Co-Morbidities. J Alzheimers Dis, doi:10.3233/JAD-190241 (2019).

42 Metaxakis, A. et al. Lowered insulin signalling ameliorates age-related sleep fragmentation in Drosophila. PLoS Biol 12, e1001824 (2014).

43 Chemelli, R. M. et al. Narcolepsy in orexin knockout mice: molecular genetics of sleep regulation. Cell 98, 437–451 (1999).

44 Nishino, S., Ripley, B., Overeem, S., Lammers, G. J. & Mignot, E. Hypocretin (orexin) deficiency in human narcolepsy. Lancet 355, 39–40, doi:10.1016/S0140-6736(99)05582-8 (2000).

45 Hara, J. et al. Genetic ablation of orexin neurons in mice results in narcolepsy, hypophagia, and obesity. Neuron 30, 345–354 (2001).

46 Bastianini, S., Silvani, A., Berteotti, C., Lo Martire, V. & Zoccoli, G. High-amplitude theta wave bursts during REM sleep and cataplexy in hypocretin-deficient narcoleptic mice. J Sleep Res 21, 185–188, doi:10.1111/j.1365-2869.2011.00945.x (2012).

47 Lee, M. G., Hassani, O. K. & Jones, B. E. Discharge of identified orexin/hypocretin neurons across the sleep-waking cycle. J Neurosci 25, 6716–6720 (2005).

48 Mileykovskiy, B. Y., Kiyashchenko, L. I. & Siegel, J. M. Behavioral Correlates of Activity in Identified Hypocretin/Orexin Neurons. Neuron 46, 787–798, doi: https://doi.org/10.1016/j.neuron.2005.04.035 (2005).

49 Peyron, C. et al. Neurons containing hypocretin (orexin) project to multiple neuronal systems. J Neurosci 18, 9996–10015 (1998).

50 Thannickal, T. C. et al. Reduced number of hypocretin neurons in human narcolepsy. Neuron 27, 469–474 (2000).

51 Sakurai, T. & Mieda, M. Connectomics of orexin-producing neurons: interface of systems of emotion, energy homeostasis and arousal. Trends Pharmacol Sci 32, 451–462, doi:10.1016/j.tips.2011.03.007 (2011).

52 Mander, B. A., Winer, J. R. & Walker, M. P. Sleep and Human Aging. Neuron 94, 19–36, doi:10.1016/j.neuron.2017.02.004 (2017).

53 Breese, C. R., Ingram, R. L. & Sonntag, W. E. Influence of age and long-term dietary restriction on plasma insulin-like growth factor-1 (IGF-1), IGF-1 gene expression, and IGF-1 binding proteins. J. Gerontol 46, B180–B187 (1991).

54 Muller, A. P. et al. Reduced brain insulin-like growth factor I function during aging. Mol Cell Neurosci 49, 9–12 (2012).

55 Kessler, B. A., Stanley, E. M., Frederick-Duus, D. & Fadel, J. Age-related loss of orexin/hypocretin neurons. Neuroscience 178, 82–88 (2011).

56 Cipriani, G., Lucetti, C., Danti, S. & Nuti, A. Sleep disturbances and dementia. Psychogeriatrics 15, 65–74, doi:10.1111/psyg.12069 (2015).

57 Ju, Y. E., Lucey, B. P. & Holtzman, D. M. Sleep and Alzheimer disease pathology--a bidirectional relationship. Nat Rev Neurol 10, 115–119, doi:10.1038/nrneurol.2013.269 (2014).

58 Swaab, D. F., Fliers, E. & Partiman, T. S. The suprachiasmatic nucleus of the human brain in relation to sex, age and senile dementia. Brain Res 342, 37–44, doi:10.1016/0006-8993(85)91350-2 (1985).

59 van Oostrom, S. H., Nooyens, A. C. J., van Boxtel, M. P. J. & Verschuren, W. M. M. Long sleep duration is associated with lower cognitive function among middle-age adults - the Doetinchem Cohort Study. Sleep Med 41, 78–85, doi:10.1016/j.sleep.2017.07.029 (2018).

60 Sindi, S. et al. Sleep disturbances and dementia risk: A multicenter study. Alzheimers Dement 14, 1235–1242, doi:10.1016/j.jalz.2018.05.012 (2018).

61 Hung, C. M. et al. Risk of dementia in patients with primary insomnia: a nationwide population-based case-control study. BMC Psychiatry 18, 38, doi:10.1186/s12888-018-1623-0 (2018).

62 Larsson, S. C. & Wolk, A. The Role of Lifestyle Factors and Sleep Duration for Late-Onset Dementia: A Cohort Study. J Alzheimers Dis 66, 579–586, doi:10.3233/JAD-180529 (2018).

63 Chen, J. C. et al. Sleep duration, cognitive decline, and dementia risk in older women. Alzheimers Dement 12, 21–33, doi:10.1016/j.jalz.2015.03.004 (2016).

64 Carro, E. & Torres-Aleman, I. The role of insulin and insulin-like growth factor I in the molecular and cellular mechanisms underlying the pathology of Alzheimer’s disease. Eur. J Pharmacol 490, 127–133 (2004).

65 Talbot, K. et al. Demonstrated brain insulin resistance in Alzheimer’s disease patients is associated with IGF-1 resistance, IRS-1 dysregulation, and cognitive decline. J. Clin. Invest 122, 1316–1338 (2012).

66 Date, Y. et al. Distribution of orexin/hypocretin in the rat median eminence and pituitary. Molecular Brain Research 76, 1–6, doi: https://doi.org/10.1016/S0169-328X(99)00317-4 (2000).

