## Supplementary figures and images for "Insulin-Like Growth Factor I Modulates Sleep Through Hypothalamic Orexin Neurons"

### Supplemental Figure 1

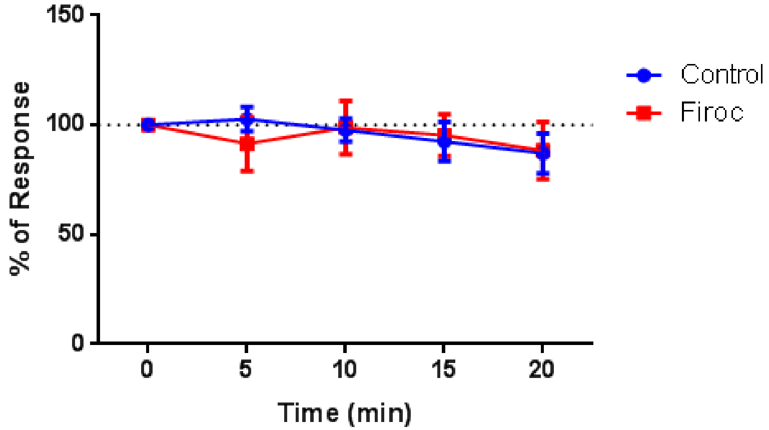


Supplementary Figure
